# DeSCENT: Deconvolutional Single-Cell RNA-seq Enhances Transcriptome-based Cancer Survival Analysis

**DOI:** 10.64898/2026.03.15.711877

**Authors:** Yonghao Zhao, Zeyu You, Yu Shen, Jielei Chu, Xun Gong, Tianrui Li, Ziqiang Wang, Chuan Xu, Zhipeng Luo, Yazhou He

## Abstract

**Motivation:** Accurate cancer survival prediction requires modeling tumor heterogeneity across both population and cell levels. Most cancer survival analyses use tumor transcriptomes only, since cohorts are usually measured with bulk RNA-seq but are rarely recorded with single-cell RNA-seq. This prevents the direct use of cell-level transcriptomes in cancer survival analysis.

**Results:** To bridge this gap, we propose using bulk RNA-seq deconvolution algorithms to reconstruct each subject’s scRNA-seq profile from their bulk data. Then, by combining both scRNA-seq and bulk RNA-seq together with their survival labels (paired to bulk), we perform multimodal transcriptome-based survival analysis. We built this framework as DeSCENT and evaluated it with common survival models on eight TCGA cancer cohorts. Results showed notable and consistent improvements in C-index over bulk-only models or models using cellular information alone.

**Availability:** Our code is available at GitHub: https://github.com/YonghaoZhao722/DeSCENT.

**Supplementary information:** Supplementary data are available at Bioinformatics online.

## 1. Introduction

Cancer remains a major global public health challenge and continues to be a leading cause of human deaths worldwide Siegel et al. (2025). It is a complex disease driven by environmental factors and genetic mutations, which together lead to malignant transformation and progression Martínez-Jiménez et al. (2020); Hanahan (2022). In the medical field, cancer prognosis and survival analysis are important feedback for precision medicine. However, performing accurate survival analysis can be quite challenging, for the cancer outcome is often affected by numerous factors such as a patient’s clinical features, pathological conditions, and tumor transcriptomes (RNA sequencing, or RNA-seq) . While plenty of work has been devoted to studying the first two factors, transcriptome-based survival analysis is still underexplored (especially on the *cell level*), primarily because the data foundation is flawed. On one hand, public databases such as TCGA The Cancer Genome Atlas Research Network (2013) and GEO Barrett et al. (2012) have offered many cancer cohorts that have paired bulk RNA-seq and survival labels, motivating numerous survival analyses that use tumor-level transcriptomes Gross et al. (2024). On the other hand, cohorts are rarely recorded with both cell-level RNA-seq and survival labels. Such a situation greatly hinders the discovery of cell-level RNA-seq that can be vital to cancer prognosis.

Recently, single-cell RNA sequencing (scRNA-seq) results have gradually become available. The improved resolution of cellular composition and states offers new perspectives for studying tumor heterogeneity. However, scRNA-seq data have limited use due to their high costs and small sample sizes. As a result, scRNA-seq data are rarely paired with survival labels, preventing their direct use in survival analysis. But fortunately, the recently proposed bulk RNA-seq deconvolution methods have shown great potential in bridging the gap in data. Bulk RNA-seq deconvolution aims to infer cell-type proportions from bulk RNA-seq according to known single-cell gene expression profiles. Examplar work includes Scaden Menden et al. (2020), TAPE Chen et al. (2022b), and ReDeconv Lu et al. (2025). Furthermore, with the help of generative models such as scVI Lopez et al. (2018), scGAN Marouf et al. (2020), and scDiffusion Luo et al. (2024), scRNA-seq profiles of different cell types can be accurately synthesized. Such advances inspire us to explore cancer survival analysis that can use cell-level transcriptomes.

We thus propose and develop a new framework named **DeSCENT** (**De**convolutional **S**ingle-**C**ell RNA **EN**hances **T**ranscriptome-based cancer survival analysis). The essential idea is to employ bulk RNA-seq deconvolution algorithms to complete one’s scRNA-seq based on their bulk; then, by combining both sc- and bulk RNA-seq, along with their survival labels that pair with bulk, we perform a full-scale transcriptome-based survival analysis. Specifically, we first apply ReDeconv Lu et al. (2025), a SOTA deconvolution method, to estimate cell type proportions from one’s bulk RNA-seq; then, by using a pretrained diffusion model similar to scDiffusion Luo et al. (2024), we can generate matching scRNA-seq data. Treating bulk RNA-seq and its generated scRNA-seq as paired modalities, we devise a feature extraction and modal fusion module that can produce complementary multi-scale transcriptome features. The key techniques used in this module are bulk-anchored contrastive alignment, gene-level mask reconstruction, and cross-attention mechanism. Finally, the transcriptome features and the original survival labels can be fed for training any survival models.

To the best of our knowledge, our work is pioneering transcriptome-based cancer survival analysis that uses both bulk RNA-seq and (deconvolved) scRNA-seq data. Our main contributions are summarized as follows.

- We leverage bulk RNA-seq deconvolution methods to generate cell-level transcriptomic profiles that have paired survival labels, overcoming the data gap present in transcriptome-based survival analysis.
- Working with RNA-seq of different resolutions, we develop a feature extraction and modal fusion module that can produce complementary multi-scale transcriptome features. These features can universally work with any survival models.
- We systematically tested DeSCENT’s various implementations working with common survival models on eight TCGA cancer cohorts. Results demonstrated notable and consistent improvements w.r.t. C-index compared to bulk RNA-seq-only models or those that use cellular information alone. Hence, DeSCENT’s efficacy and robustness have been fully validated.

## 2. Related Work

### 2.1. Bulk RNA-seq Deconvolution

A tumor’s gene expression profiles are measured as bulk RNA-seq, which is an arithmetic sum of single-cell RNA-seq of many cell types. Due to this relation, bulk RNA-seq can be deconvolved into cell-type proportions, given that proper cell-type gene expression profiles are referred to. Classic deconvolution methods, such as CIBERSORTx Newman et al. (2019), MuSiC Wang et al. (2019), and BayesPrism Chu et al. (2022), use regression or Bayesian models to estimate cell fractions; others like Scaden Menden et al. (2020) and TAPE Chen et al. (2022b) employ neural nets to realize tissue-adaptive deconvolution and cell-type-specific expression prediction. Notably, the latest work, ReDeconv Lu et al. (2025), proposed a normalization technique called CLTS (Count based on Linearized Transcriptome Size), which eliminated sample biases resulting from different transcriptome sizes and sequencing depths.

As mentioned earlier, existing scRNA-seq data are relatively limited in scale and paired annotations. So, people are exploring generative models to synthesize single-cell expression profiles. Early examples are scGAN Marouf et al. (2020), scDesign3 Song et al. (2024), and scVI Lopez et al. (2018), which use GANs, probabilistic generative models, and VAEs, respectively. More recently, diffusion-based and flow-based models have shown stronger generation performance and are being adopted in many single-cell analyses. scDiffusion Luo et al. (2024) couples pre-trained foundation models with conditional diffusion to generate multi-condition single-cell data; CFGen Palma et al. (2025) uses a conditional flow matching algorithm plus discrete likelihood decoders to generate multimodal single-cell counts; and scRDiT Dong et al. (2025) adopts a diffusion-transformer for performing high-fidelity scRNA-seq synthesis.

### 2.2. Multimodal Cancer Survival Analysis

Richer oncological data resources are motivating multimodal cancer survival analysis, which can comprehensively characterize tumor heterogeneity and its impact on cancer prognosis. To date, common modalities employed in cancer survival analysis include clinical features, pathological whole-slide images (WSIs), and bulk transcriptomes. Representative works are Pathomic Fusion Chen et al. (2022a), which uses tensor fusion to jointly model histopathological and genetic features, and POMP Wang et al. (2025), which aligns WSIs and multi-omics representations in a shared latent space and achieved superior performance on multiple TCGA cohorts. It is worth mentioning that the multi-omics used in POMP are all bulk-level modalities, thus not explicitly modeling the effects of cell-level omics.

There are indirect uses of scRNA-seq plus bulk RNA-seq in cancer survival analysis. Their typical pattern is to use scRNA-seq only in upstream tasks such as gene selection, and the final survival model still operates on bulk data alone. For example, Cai et al. Cai et al. (2025) uses scRNA-seq to identify prognostic genes, while scBGDL Liu et al. (2025) uses scRNA-seq to construct gene–gene relational graphs. More recently, **scSurv** Mizukoshi et al. (2025) is the first study that uses cellular transcriptome information directly in survival analysis. They first deconvolute bulk RNA-seq into cell-state–level mixture weights and then use them in modeling patient risk via a Cox formulation. This design aims at relating survival risk to cell states and has offered enhanced interpretability at the cell level. But because scSurv only focuses on cells, the survival prediction performance it achieves is inferior (see our experiments below). By comparison, our work uses bulk and (deconvolved) single-cell RNA-seq together as paired input modalities in survival analysis. The completed scRNA-seq provides complementary, fine-grained cellular information to its bulk counterpart, thus jointly improving the survival prediction.

## 3. Methods

Our proposed framework, DeSCENT, aims to perform transcritome-based cancer survival analysis that uses both bulk and single-cell RNA-seq. It consists of a deconvolution module and a multimodal representation module (Fig. 1). The objective of the first one is to construct a survival cohort where each patient is associated with both bulk and single-cell RNA-seq (scRNA-seq) data. We particularly employed ReDeconv Lu et al. (2025), a SOTA deconvolution method, and a pretrained diffusion model to generate very accurate scRNA-seq from its bulk. Secondly, we fuse bulk and single-cell RNA-seq data into aligned features, which are fed for training survival models. The rest of this section details DeSCENT’s design and implementation.

**Figure 1.**
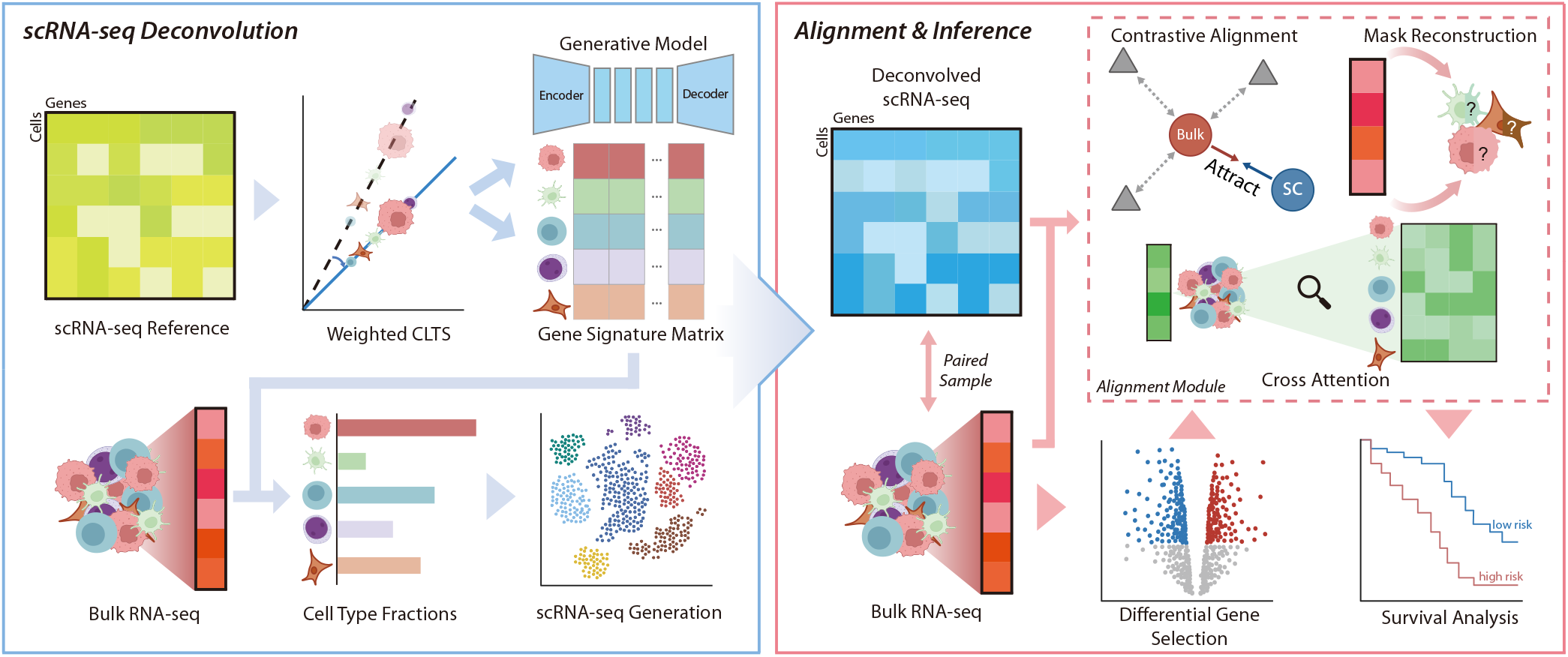
Overview of DeSCENT. (Left) The scRNA-seq deconvolution module. An external scRNA-seq dataset, after properly normalized by ReDeconv’s CLTS technique, serves as the reference for estimating the cell-type fractions of one’s bulk RNA-seq; conditioning on these fractions, scDiffusion is used to generate patient-specific scRNA-seq. (Right) Multimodal representation module. One’s bulk and scRNA-seq are aligned via contrastive learning and mask reconstruction techniques, and are fused by a cross-attention mechanism. The learned representations are fed to a downstream survival model. Created with BioRender.com.

### 3.1. Data Preprocessing

#### Preliminaries

The primary data needed for performing transcriptome-based cancer survival analysis is a cancer cohort 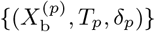, where each patient *p* is associated with a bulk expression vector 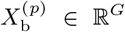 in TPM (transcripts per million), an observed survival time *T*_*p*_, and a censoring indicator *δ*_*p*_ ∈ {0, 1}. Such datasets are publicly available, such as from TCGA. Our first goal is to generate additional scRNA-seq 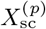 by deconvolution according to their bulk 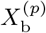. Here 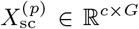 is a matrix in UMI (unique molecule identifier), *c* the total number of cells, and *G* the total number of genes. Hence, we will have a richer dataset 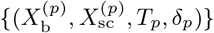 to do survival. To realize deconvolution, we also need another scRNA-seq reference dataset *D*_ref_ of the same cancer type. *D*_ref_ is required to be annotated with cell types and serves as a biological prior for cell-type–specific expression patterns.

#### Reference data normalization

We normalize the scRNA-seq reference *D*_ref_ using *Weighted CLTS*, an extension of ReDeconv’s CLTS, to reduce biases induced by heterogeneous transcriptome sizes across subjects and batches. Assume *D*_ref_ contains *S* samples and *K* annotated cell types. Each reference cell *i* has a cell-type label *k*(*i*) ∈ {1, …, *K*} and belongs to a sample *s*(*i*) ∈ {1, …, *S*}. Let *x*_*ij*_ denote the raw UMI count of gene *j* in cell *i* and define the transcriptome size (total UMI) of cell *i* as

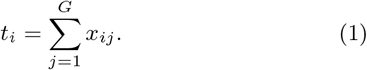

For each sample *m* and cell type *k*, denote the corresponding cell set and its size by

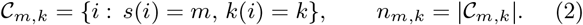

When *n*_*m,k*_ *>* 0, the sample-wise, cell-type-wise mean transcriptome size is

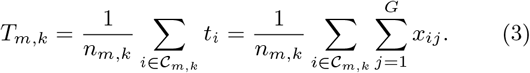

Following CLTS, we choose a baseline sample (without loss of generality, *m* = 1) and estimate, for each sample *m*, a multiplicative size factor *a*_*m*_ *>* 0 that aligns the cell-type mean transcriptome sizes of sample *m* to the baseline. Let 𝒦_*m*_ = {*k* : *n*_*m,k*_ *>* 0 ∧ *n*_1,*k*_ *>* 0} be the set of cell types shared by sample *m* and the baseline. Under the no-shift setting, *a*_*m*_ is obtained by least squares through the origin:

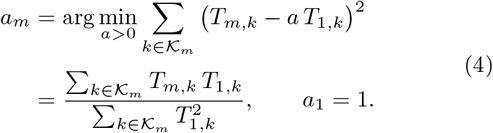

All cells in sample *m* are then rescaled by

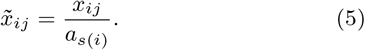

To avoid outsized influence from extremely rare cell types, we use an abundance-weighted variant. Define

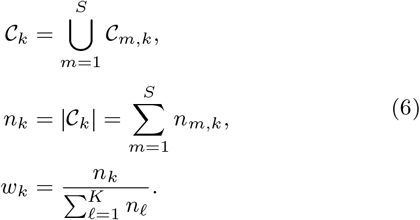

With 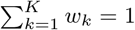. Weighted CLTS estimates

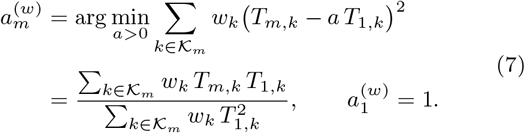

and applies the same no-shift rescaling:

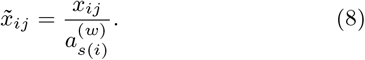

When *w*_*k*_ is uniform over 𝒦_*m*_, Weighted CLTS reduces to the unweighted CLTS. In practice, we compute {*T*_*m,k*_} on raw UMI counts in *D*_ref_, estimate {*w*_*k*_} from annotated cell-type frequencies, and normalize all reference cells before constructing the signature matrix for deconvolution.

#### DEG selection

For each cancer type, we restrict our deconvolution to a list of genes that is shared by the bulk RNA-seq cohort and the scRNA-seq reference data. This preserves around 15k to 29k genes depending on different cancers. When it comes to the multimodal fusion and survival analysis module, a subset of genes is further selected by doing differential expression analysis. We adopt PyDESeq2 Muzellec et al. (2023), which selects differentially expressed genes (DEGs) on the bulk RNA-seq cohort by comparing cancer samples versus normal samples using only the training subset in each split. Genes with Benjamini-Hochberg adjusted *p*-value *<* 0.05 and | log_2_ FoldChange| *>* 1.5 are retained. This results in 2k to 4k DEGs selected. See supplementary (S1) for the statistics.

### 3.2. scRNA-seq Generation

Our first module is to generate 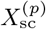 based on its bulk 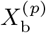 for each patient *p*. This is done in two steps: the first step is, according to the reference data *D*_ref_, to estimate (or deconvolute) the cell-type fractions that make up a bulk; the second step is to generate realistic scRNA-seq by a generative model conditioned on the cell-type fractions.

#### Bulk deconvolution

We first apply ReDeconv for estimating the cell-type fractions of each bulk sample. ReDeconv links bulk expression to cell-type–specific signatures learned from the single-cell reference *D*_ref_ and outputs a patient-level cell-type fraction vector:

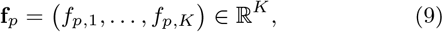

where *K* is the number cell types, and *f*_*p,k*_ ∈ [0, 1] is a fraction of cell type *k* with 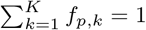.

#### scRNA-seq generation

In order to provide a biologically grounded prior over cell-type–specific gene expression patterns, we adopt scDiffusion Luo et al. (2024), a conditional latent diffusion model trained solely on the scRNA-seq reference *D*_ref_ . Then, we can generate patient-specific single-cell data 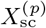 conditioned on **f**_*p*_ by the trained scDiffusion model. Specifically, we fix a global total number of generated cells *c* per patient and allocate *c*_*p,k*_ = ⌊*c* · *f*_*p,k*_⌋ cells to each cell type *k*. We then sample *c*_*p,k*_ cells from scDiffusion conditioned on cell type *k*, yielding a deconvolved single-cell expression matrix

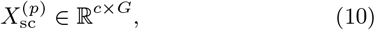

where *G* is the total number of genes used in deconvolution.

#### Remarks

We want to stress that mapping bulk RNA-seq to its cell-level compositions is inherently non-invertible, thus being an ill-posed problem. Accordingly, DeSCENT does not aim to infer a ground-truth single-cell realization. Instead, we step back, only attempting to offer an auxiliary cellular modality (constrained by the bulk) necessary for performing certain downstream tasks (i.e., survival analysis) Koo et al. (2025).

### 3.3. Multimodal Representation

Now we have a multiscale dataset 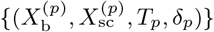 prepared, with 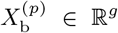 being a patient’s bulk RNA-seq vector, 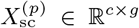 the generated scRNA-seq matrix, and (*T*_*p*_, *δ*_*p*_) the survival labels. Note that, for the rest of this section, *g* denotes the number of differentially expressed genes (DEGs), i.e., a selected subset of the original *G* genes. The second module here (Fig. 2) is to co-learn the representations of *X*_b_ and *X*_sc_ (sometimes the superscript *p* is dropped if it causes no confusion) by contrastive learning and mask reconstruction, and then fuse them by cross-attention. The fused representation can be prelearned or co-learned with the downstream survival models.

**Figure 2.**
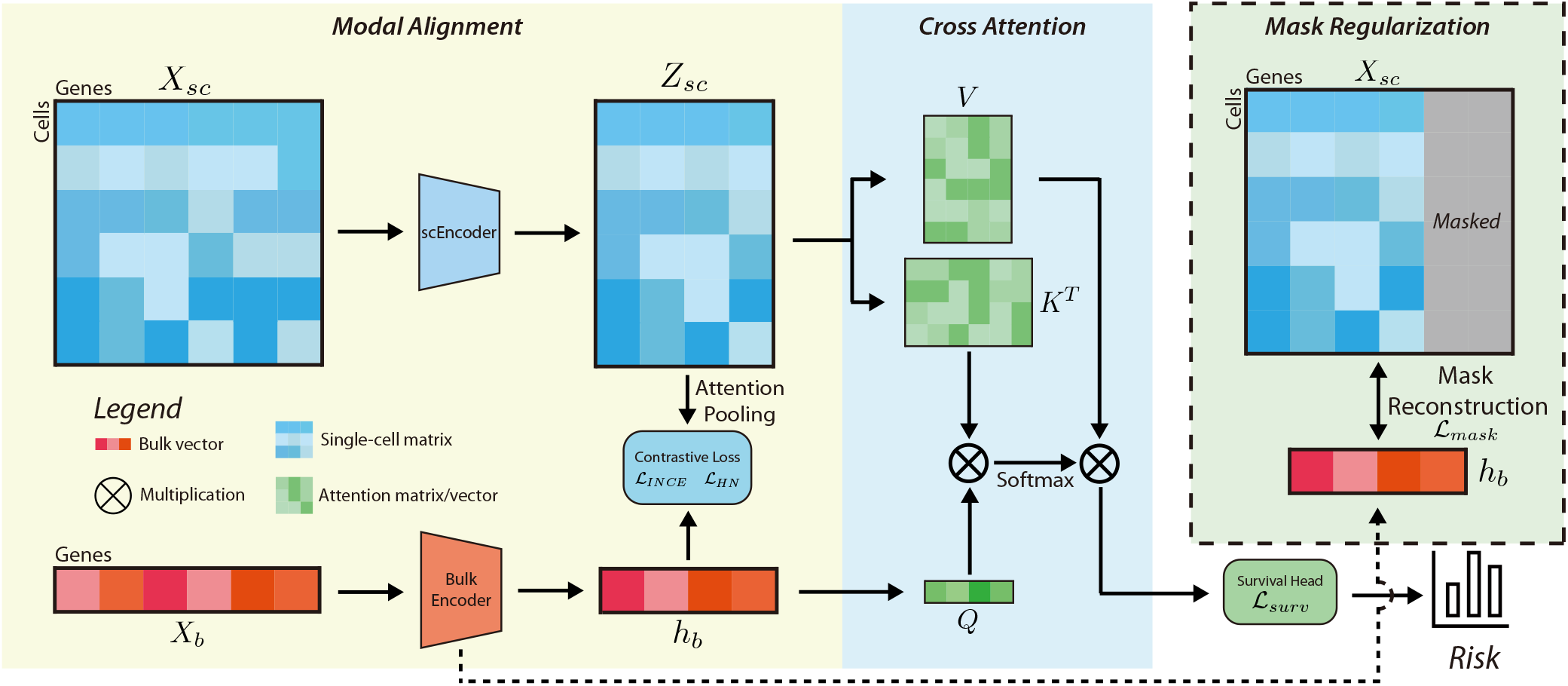
Architecture of the multimodal fusion module, i.e. the right-hand-side module in Fig. 1. First, each patient’s scRNA-seq matrix *X*_sc_ and its bulk RNA-seq vector *X*_b_ are aligned in a latent space. This is done by encoding *X*_sc_, *X*_b_ to lower-dimensional representations *Z*_sc_, *h*_b_, respectively, which are further aligned by contrastive learning. In parallel, *h*_b_ is jointly optimized with a mask-reconstruction head to reconstruct masked entries of *X*_sc_. Lastly, to do multimodal fusion, *Z*_sc_ is projected to a key-value pair (*K, V*) and *h*_b_ to a query *Q*, which are fused by cross-attention that produces a combined representation fed for a downstream survival model.

#### Latent representations

We first map *X*_sc_ to cell-level embeddings using a single-cell encoder *f*_sc_, consisting of a feed-forward embedding network followed by multi-head self-attention (MHSA) across cells. LayerNorm is applied at each layer and on the final output to stabilize training and improve representation robustness. This yields context-enriched cell representations

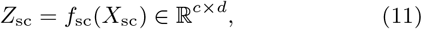

where *d* denotes the latent dimension. To obtain a patient-level single-cell representation, we further apply an attention pooling module to the contextualized cell embeddings,

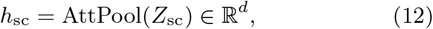

where the learned attention weights reflect the relative importance of individual cells. In parallel, the bulk vector *X*_b_ is passed through a bulk encoder *f*_bulk_, which follows the same design as in the bulk-only baseline for fair comparison, to learn a bulk-level embedding

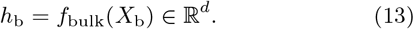

#### Contrastive alignment

To align bulk and single-cell embeddings at the patient level, we adopt two contrastive learning losses that were commonly used van den Oord et al. (2018); Chen et al. (2020). The first one is an InfoNCE loss ℒ_INCE_ that treats each bulk embedding as an anchor, its paired single-cell embedding as positive, and others negative. The rationale of using bulk embedding as the anchor since that bulk expressions are directly observed, thus being a more reliable reference. To be computationally efficient, we do contrastive losses in minibatches. Concretely, denote by 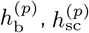 the bulk and single-cell embeddings of a patient *p* after a shared *ℓ*_2_-normalized projection. For a minibatch ℬ in stochastic training, we define a similarity score sim 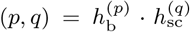. Then, a single-direction InfoNCE loss is defined as:

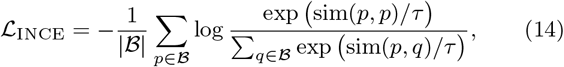

where *τ* is a temperature hyperparameter.

We further incorporate a hard-negative matching loss ℒ_HN_ that focuses on the most confusable negative point under the same anchor direction. That is, the hardest negative index for an anchor *p* is defined as

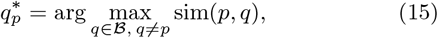

leading to a two-class softmax loss

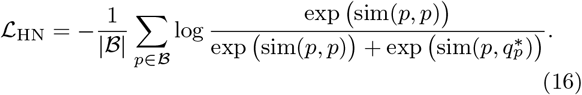

The two losses are weighted separately but are jointly optimized during training.

#### Mask reconstruction

In parallel, to enhance the coupling between bulk and single-cell embeddings, we introduce a self-supervised mask reconstruction loss. At each training epoch, we randomly mask 20% of the gene entries in *X*_sc_ to obtain a partially observed matrix 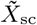. Let **M** ∈ {0, 1}^*c×g*^ denote the corresponding binary mask, where **M**_*ij*_ = 1 indicates a masked gene entry. The bulk latent vector *h*_b_ is concatenated with 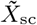 to form the reconstruction input. Reconstructions are done by a two-layer reconstruction head with input dimension being *g* + *d* and output dimension *g*, producing a reconstructed matrix 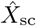. We define the mask reconstruction loss as

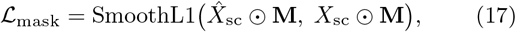

where SmoothL1(·, ·) is the smooth-*ℓ*_1_ loss, and ⊙ denotes element-wise multiplication. The loss is evaluated only on masked positions.

#### Representation fusion

To fuse bulk and single-cell embeddings, we apply a cross-attention mechanism between *h*_b_ and *Z*_sc_. We first map *h*_b_ to a query vector *Q*, and *Z*_sc_ to key and value matrices *K* and *V* :

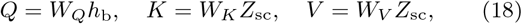

where *W*_*Q*_, *W*_*K*_, *W*_*V*_ are projection matrices of attention heads. Then the cross-attention output is given by

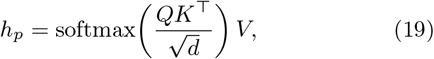

which aggregates cell-level information into a fused patient representation for survival modeling.

#### Survival head

The fused embedding *h*_*p*_ is used as covariates in a survival head for risk prediction. We explore three types of survival heads: (i) a Cox model, (ii) a DeepSurv model with an MLP on top of *h*_*p*_, and (iii) a discrete-time DeepHit-style model Katzman et al. (2018); Lee et al. (2018). For the Cox and DeepSurv heads, we obtain a scalar risk score *r*_*p*_ and optimize the negative partial log-likelihood,

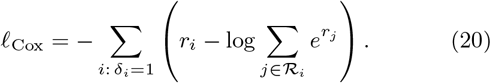

We further apply elastic net regularization to the Cox layer parameters, leading to the final loss

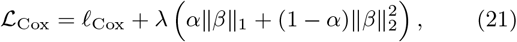

where *δ*_*i*_ is the event indicator and ℛ_*i*_ is the risk set. DeepSurv differs only by using an MLP to compute *r*_*p*_ but keeps the same loss form.

For the DeepHit head, we discretize time into *T* intervals and let ***π***_*p*_ ∈ ℝ^*T*^ denote the predicted event-time distribution for patient *p*, where *π*_*p,t*_ = Pr(event in interval *t* | *p*) and 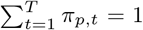. Let 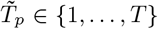 be the observed (possibly censored) interval and *δ*_*p*_ ∈ {0, 1} the event indicator. For uncensored patients (*δ*_*p*_ = 1), define one-hot event labels 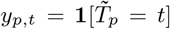. Its loss consists of a discrete log-likelihood term

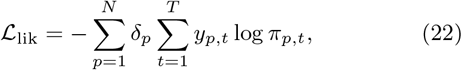

and a ranking term over censoring-aware comparable pairs,

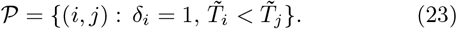

Define a scalar risk score 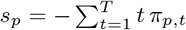, then

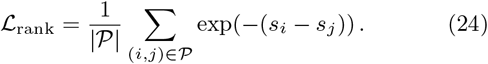

The DeepHit survival loss is ℒ_DH_ = *β*ℒ_lik_ + (1 − *β*)ℒ_rank_.

In all cases, we denote the final survival loss generically by ℒ_surv_. The total training objective is a weighted sum of the four losses,

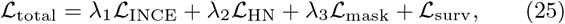

where *λ*_*i*_ *>* 0 are hyperparameters. Finally, a standard optimization procedure is used to work out the parameters.

## 4. Experiments

In this section, we evaluate DeSCENT’s performance and test whether the additionally generated cell-level transcriptomes can enhance the final survival prediction. Also, to assess DeSCENT’s universality, we apply it to three common survival models and test them on eight TCGA cancer cohorts that were frequently studied Gross et al. (2024). Key results delivered in this section are bulk deconvolution accuracy, survival performance, and ablation studies.

### 4.1. Datasets

Eight common cancer cohorts are selected from TCGA The Cancer Genome Atlas Research Network (2013): colon adenocarcinoma (COAD), breast invasive carcinoma (BRCA), lung adenocarcinoma (LUAD), hepatocellular carcinoma (LIHC), stomach adenocarcinoma (STAD), lower grade glioma (LGG), kidney renal clear cell carcinoma (KIRC), and head and neck squamous cell carcinoma (HNSC). To enable deconvolution, we curated a matched scRNA-seq reference dataset for each cancer type from publicly available resources: COAD: HTAN VUMC Chen et al. (2021), BRCA: GSE176078 Wu et al. (2021), LUAD: GSE131907 Kim et al. (2020), LIHC: GSE149614 Lu et al. (2022), STAD: GSE183904 Kumar et al. (2022), LGG: GSE232316 Zahedi et al. (2025), KIRC: GSE210042 Davidson et al. (2023), HNSC: GSE181919 Choi et al. (2023), As the STAD and KIRC scRNA-seq datasets lack cell-type annotations, we employ sc-type Ianevski et al. (2022) to perform automated annotations based on marker genes. Data details such as the number of cohorts and genes are reported in the supplementary (S1).

#### Data preprocessing

For each cancer type, we first intersect the Gene IDs between the bulk RNA-seq data and the corresponding scRNA-seq reference. All subsequent analyses are performed only on this shared gene list. The scRNA-seq expression matrices and metadata are formatted uniformly, followed by the weighted CLTS normalization.

### 4.2. Results

#### Bulk deconvolution

Foremost, we must ensure that the generated scRNA-seq are accurate enough. As paired bulk RNA-seq and scRNA-seq datasets are rare in practice, prior work often uses pseudo (bulk, sc) RNA-seq samples for evaluation. To do so, we split each scRNA-seq reference dataset into two (4:1), the former used for training scDiffusion and the rest for constructing pseudo samples. To create a (bulk, sc) pair, an scRNA-seq sample is randomly sampled from the constructing data and is then aggregated into a bulk sample. The cell-type fractions of these pseudo samples are thus known and serve as the ground truth. To perform deconvolution, we test three deconvolution models: ReDeconv, TAPE, and Scaden, which estimate cell-type fractions of a pseudo bulk. Then, these fractions guide the trained scDiffusion to generate scRNA-seq, which are compared against the corresponding pseudo scRNA-seq. The metrics used for comparison are the Pearson correlation, Spearman correlation, and Maximum Mean Discrepancy (MMD) Gretton et al. (2012). As shown in Supplementary Section, ReDeconv achieves the best average performance and is the top method on almost all cohort–metric pairs.

#### Survival analysis

We next evaluate the performance of DeSCENT’s survival predictions. The base survival models used are Cox and two neural network based models, DeepSurv Katzman et al. (2018) and DeepHit Lee et al. (2018). To demonstrate the improvement contributed by the additional scRNA-seq data, we compare with three baselines, `Bulk`, `Bulk+CT`, and `scSurv` Mizukoshi et al. (2025). The first one only uses bulk RNA-seq as input features, and the second one additionally uses cell type proportions (CT) given by ReDeconv. Specifically, `Bulk+CT` first keeps the bulk encoder and survival head identical to `Bulk` for a fair comparison; meanwhile, a lightweight cell-type branch is added, which encodes the *K*-dimensional cell-type fractions through a small MLP to a 256-dimensional embedding that is fused with the bulk representation. The third one is aforementioned `scSurv`, which uses cellular transcriptome information alone to perform survival. Note that to the best of our knowledge, there are no comparable survival baselines that take patient-level bulk RNA-seq and paired scRNA-seq modalities (even synthesized) as inputs under the same task setting. More implementation details of the above methods and ours can be found in Supplementary Section.

Table 1 reports the absolute C-index values that evaluate all models’ survival performance on all cancer cohorts. For each cohort, we perform five independent 65/15/20 train/validation/test splits. In each split, the 20% test set is fully held out and never used for feature selection or model fitting, and reported values are averages over these five held-out tests. Firstly, we stress that `Bulk`’s results worked out by us are consistent with the latest large-scale benchmark test offered by Gross et al. (2024). They similarly tested penalized Cox, DeepSurv, and DeepHit on numerous TCGA cancer cohorts using bulk RNA-seq alone under standard feature selection and held-out evaluation settings. Second, we see that `Bulk+CT` performs comparably to, or slightly worse than `Bulk`, indicating that a naive representation of cell-type vectors fused to their bulk fails to provide sufficient complementary prognostic signals. Thirdly, `scSurv` performs very poorly on the survival prediction, although it enjoys some extent of interpretability. In contrast, DeSCENT achieves consistently superior performance across almost all cancer types regardless of the base survival models. In detail, compared to the averaged C-index (across eight cancers) achieved by `Bulk`, DeSCENT-Cox improves by 9.1%, DeSCENT-DeepSurv by 6.4%, and DeSCENT-DeepHit by 3.7%. For each cancer, the best C-index attained by DeSCENT reaches 0.708 on COAD, 0.729 on BRCA, 0.853 on LGG, 0.731 on LIHC, 0.702 on LUAD, 0.627 on STAD, 0.743 on KIRC, and 0.664 on HNSC. Based on the improvements made by DeSCENT, we can draw two conclusions: 1) the generated scRNA-seq data does enhance the survival prediction, and 2) combining both sc- and bulk RNA-seq can achieve the top performance.

**Table 1.**
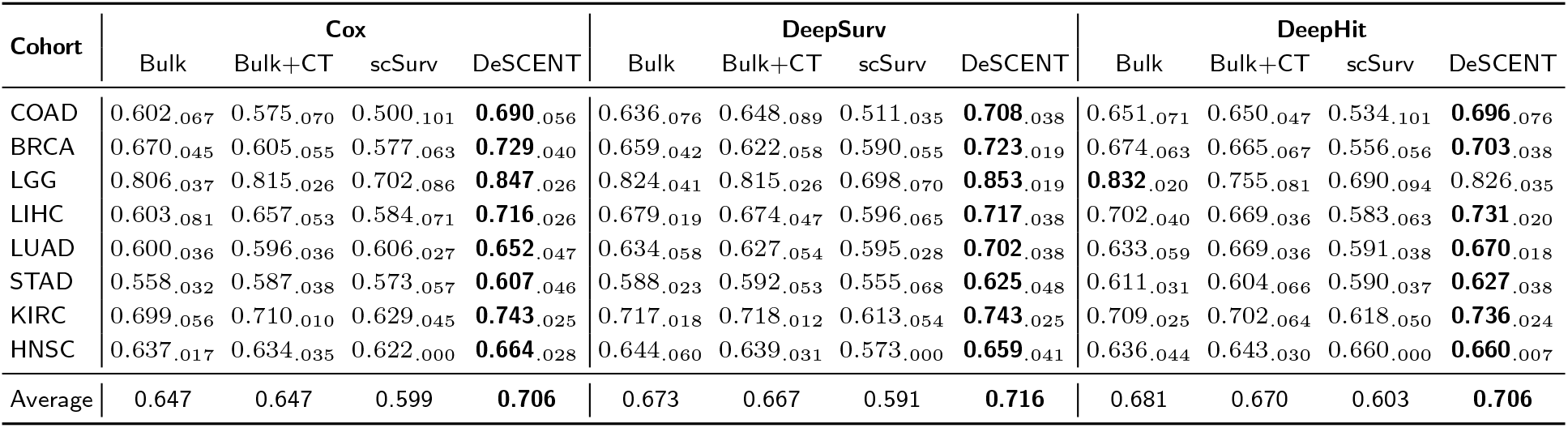
Performance comparison of different methods under three survival models across eight cancer cohorts. Averaged C-index over five held-out test runs is reported, with the standard deviation shown as a subscript. Bold indicates the best method within each survival model for each cohort.

| Cohort | Cox |  |  |  | DeepSurv |  |  |  | DeepHit |  |  |  |
| --- | --- | --- | --- | --- | --- | --- | --- | --- | --- | --- | --- | --- |
|  | Bulk | Bulk+CT | scSurv | DeSCENT | Bulk | Bulk+CT | scSurv | DeSCENT | Bulk | Bulk+CT | scSurv | DeSCENT |
| COAD | 0.602<sub>.067</sub> | 0.575<sub>.070</sub> | 0.500<sub>.101</sub> | **0.690**<sub>.056</sub> | 0.636<sub>.076</sub> | 0.648<sub>.089</sub> | 0.511<sub>.035</sub> | **0.708**<sub>.038</sub> | 0.651<sub>.071</sub> | 0.650<sub>.047</sub> | 0.534<sub>.101</sub> | **0.696**<sub>.076</sub> |
| BRCA | 0.670<sub>.045</sub> | 0.605<sub>.055</sub> | 0.577<sub>.063</sub> | **0.729**<sub>.040</sub> | 0.659<sub>.042</sub> | 0.622<sub>.058</sub> | 0.590<sub>.055</sub> | **0.723**<sub>.019</sub> | 0.674<sub>.063</sub> | 0.665<sub>.067</sub> | 0.556<sub>.056</sub> | **0.703**<sub>.038</sub> |
| LGG | 0.806<sub>.037</sub> | 0.815<sub>.026</sub> | 0.702<sub>.086</sub> | **0.847**<sub>.026</sub> | 0.824<sub>.041</sub> | 0.815<sub>.026</sub> | 0.698<sub>.070</sub> | **0.853**<sub>.019</sub> | **0.832**<sub>.020</sub> | 0.755<sub>.081</sub> | 0.690<sub>.094</sub> | 0.826<sub>.035</sub> |
| LIHC | 0.603<sub>.081</sub> | 0.657<sub>.053</sub> | 0.584<sub>.071</sub> | **0.716**<sub>.026</sub> | 0.679<sub>.019</sub> | 0.674<sub>.047</sub> | 0.596<sub>.065</sub> | **0.717**<sub>.038</sub> | 0.702<sub>.040</sub> | 0.669<sub>.036</sub> | 0.583<sub>.063</sub> | **0.731**<sub>.020</sub> |
| LUAD | 0.600<sub>.036</sub> | 0.596<sub>.036</sub> | 0.606<sub>.027</sub> | **0.652**<sub>.047</sub> | 0.634<sub>.058</sub> | 0.627<sub>.054</sub> | 0.595<sub>.028</sub> | **0.702**<sub>.038</sub> | 0.633<sub>.059</sub> | 0.669<sub>.036</sub> | 0.591<sub>.038</sub> | **0.670**<sub>.018</sub> |
| STAD | 0.558<sub>.032</sub> | 0.587<sub>.038</sub> | 0.573<sub>.057</sub> | **0.607**<sub>.046</sub> | 0.588<sub>.023</sub> | 0.592<sub>.053</sub> | 0.555<sub>.068</sub> | **0.625**<sub>.048</sub> | 0.611<sub>.031</sub> | 0.604<sub>.066</sub> | 0.590<sub>.037</sub> | **0.627**<sub>.038</sub> |
| KIRC | 0.699<sub>.056</sub> | 0.710<sub>.010</sub> | 0.629<sub>.045</sub> | **0.743**<sub>.025</sub> | 0.717<sub>.018</sub> | 0.718<sub>.012</sub> | 0.613<sub>.054</sub> | **0.743**<sub>.025</sub> | 0.709<sub>.025</sub> | 0.702<sub>.064</sub> | 0.618<sub>.050</sub> | **0.736**<sub>.024</sub> |
| HNSC | 0.637<sub>.017</sub> | 0.634<sub>.035</sub> | 0.622<sub>.000</sub> | **0.664**<sub>.028</sub> | 0.644<sub>.060</sub> | 0.639<sub>.031</sub> | 0.573<sub>.000</sub> | **0.659**<sub>.041</sub> | 0.636<sub>.044</sub> | 0.643<sub>.030</sub> | 0.660<sub>.000</sub> | **0.660**<sub>.007</sub> |
| Average | 0.647 | 0.647 | 0.599 | **0.706** | 0.673 | 0.667 | 0.591 | **0.716** | 0.681 | 0.670 | 0.603 | **0.706** |

#### Risk stratification

To assess clinical utility, we further conduct risk stratification based on the predicted risk scores. The Kaplan-Meier curves plotted in Fig. 3 show very good separations between the low-risk and high-risk groups across all eight cancer types. The log-rank test results also confirm the statistical significance, with all the p-values below 0.05. Such results suggest that our proposed bulk + single-cell RNA-seq fusion framework not only improves C-index but also yields robust and clinically meaningful patient stratification.

**Figure 3.**
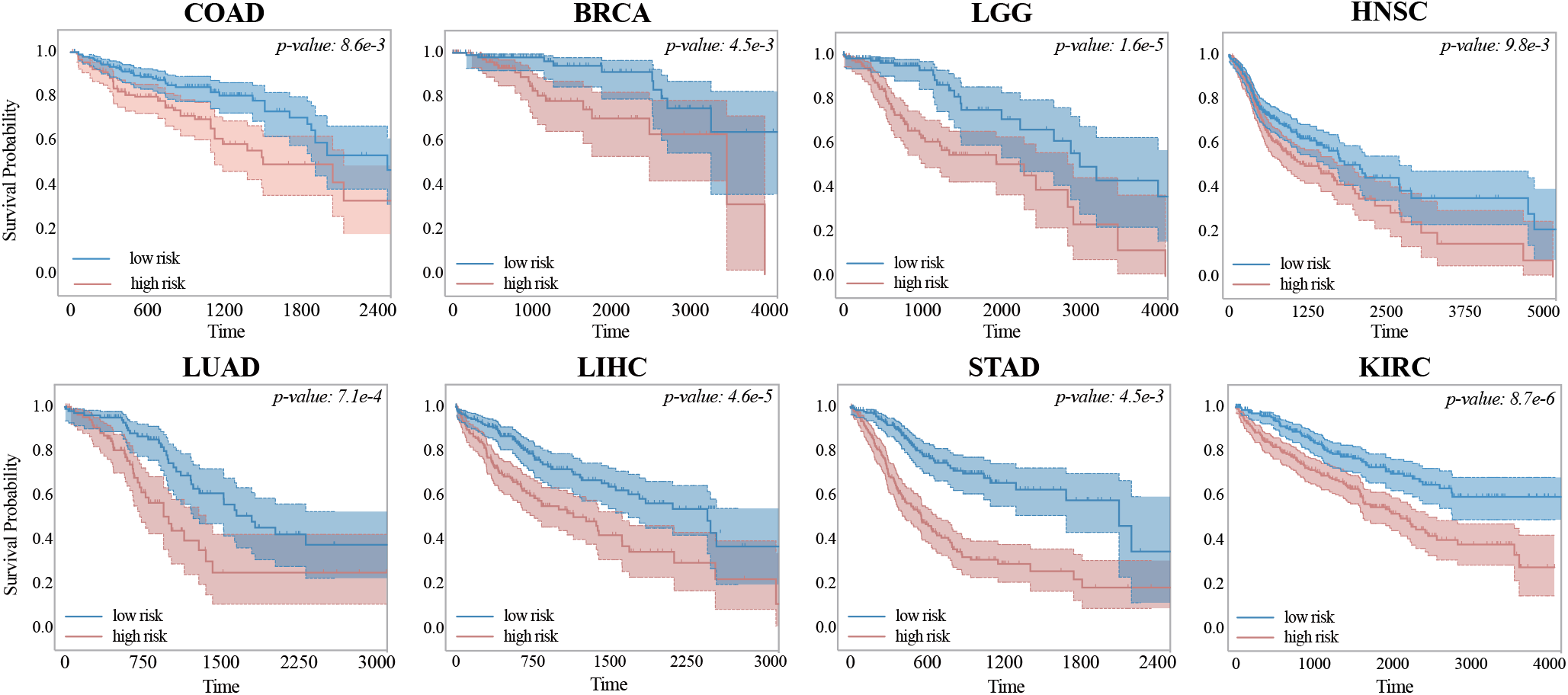
Kaplan-Meier survival analyses that assess DeSCENT. The plots demonstrate clear distinctions between the predicted low-risk and high-risk patient groups across all eight cancer cohorts. The p-values are log-rank test results.

### 4.3. Ablation Study

We also conduct an additional ablation study to investigate the contributions of different loss functions used in DeSCENT, i.e., Eq. (25). The results are shown in Supplementary Section. We can see that DeSCENT’s performance drops notably without the contrastive alignment losses (ℒ_INCE_ and ℒ_HN_, activated or deactivated together) or the mask reconstruction loss (ℒ_mask_). Moreover, we previously saw that a simple addition of deconvolved cell-type proportions (i.e. `Bulk+CT`) does not yield gains over bulk-only models. Together with the ablation results here, we can conclude that the performance gain of DeSCENT should not be attributed to cell-type proportions alone, but instead stems from modeling higher-order cellular heterogeneity through the completed single-cell modality and cross-modal fusion.

## 5. Conclusions and Future Work

We proposed and developed DeSCENT, a transcriptome-based cancer survival analysis framework using bulk seq-RNA and deconvoluted scRNA-seq data. We adopted a modified ReDeconv to perform deconvolution and designed a dual-modality representation module to align and fuse different scales of transcriptomes. Our comprehensive results are supportive of the fact that the deconvoluted scRNA-seq data, although being synthesized from their bulk, can largely enhance the survival prediction. In addition, to achieve the best prediction performance, we need to combine both sc- and bulk RNA-seq data that are complementary to each other. To effectively combine these two modalities, contrastive learning, mask reconstruction, and cross-model attention techniques are necessary functions in the framework. Lastly, having been tested with three common survival models on eight cancer cohorts, DeSCENT’s universality and robustness have been demonstrated.

We view our work as an initiative toward future cancer survival analysis using multiscale transcriptomes. With scRNA-seq data becoming more accessible and affordable, integrating bulk and single-cell transcriptomic signals is likely to evolve into an increasingly practical and powerful strategy for prognosis modeling. We believe that our work shall be a critical attempt to envision this perspective.

### Future work

Several promising directions remain for future study. First, DeSCENT provides an opportunity to extract improved interpretability and biological insights from single-cell signals. For example, analyzing the distribution of model attention over cell types and linking high-risk patterns to specific cellular components. Second, the deconvolution and modality completion process could be enhanced by incorporating richer patient-specific information, enabling more fine-grained and personalized reconstruction of single-cell profiles from bulk data.

Finally, we plan to include more modalities, such as clinical information and pathological features, into the current framework to further improve the predictive performance and push DeSCENT to clinical practices.

## Data availability

Data used in this study are available in the DeSCENT GitHub repository: https://github.com/YonghaoZhao722/DeSCENT.

