## Supplementary Material for "DeSCENT: Deconvolutional Single-Cell RNA-seq Enhances Transcriptome-based Cancer Survival Analysis"

*Bioinformatics*, YEAR, , 1–4
DOI added during production
Published: Date added during production
**Supplementary Material**

### DeSCENT: Deconvolutional Single-Cell RNA-seq Enhances Transcriptome-based Cancer Survival Analysis Supplementary Material

**Yonghao Zhao,[1,†] Zeyu You,[2,†] Yu Shen,[5] Jielei Chu,[2] Xun Gong,[2] Tianrui Li,[2] Ziqiang Wang,[5] Chuan Xu,[6] Zhipeng Luo[2,*] and Yazhou He[3,4,*]**

[1]SWJTU-Leeds Joint School, Southwest Jiaotong University, Chengdu, China
[2]School of Computing and Artificial Intelligence, Southwest Jiaotong University, Chengdu, China
[3]Yu-yue Pathology Scientific Research Center, Jinfeng Laboratory, Chongqing, China
[4]Department of Epidemiology and Biostatistics, School of Public Health, Imperial College London, London, United Kingdom
[5]Colorectal Cancer Center, Department of General Surgery, West China Hospital, Sichuan University, Chengdu, China
[6]Department of Oncology and Cancer Institute, Sichuan Academy of Medical Sciences, Sichuan Provincial People's Hospital, University of Electronic Science and Technology of China, Chengdu, Sichuan 610072, China
[†]These authors contributed equally.

#### Abstract

**Supplementary material for the manuscript DeSCENT: Deconvolutional Single-Cell RNA-seq Enhances Transcriptome-based Cancer Survival Analysis.**

### S1. Data Details

Table S1 summarizes dataset statistics for each cancer type used in our experiments, including cohort-level statistics (bulk samples and scRNA-seq donors), reference scRNA-seq annotation statistics (cells and cell types), and gene set statistics. The data sources and their roles in the pipeline are described in the main text.

### S2. Implementation Details

*Training setup.*
All experiments were implemented in PyTorch and conducted on two NVIDIA RTX 4090 GPUs. All models were optimized using AdamW with a cosine annealing learning rate schedule and a warm-up phase covering the first $5\%$ of training steps. All survival analysis models were trained and evaluated using the same DEG-restricted gene set.

*Bulk-only baseline.*
The bulk-only baseline, `Bulk`, takes the bulk RNA-seq vector as input and uses a two-layer MLP as the bulk encoder, followed by the same survival head as the corresponding method (Cox, DeepSurv, or DeepHit). The bulk encoder hidden dimensions, activation, dropout, and normalization options were tuned per cohort within the grid search space defined in Table S2.

*Bulk+CT baseline.*
The `Bulk+CT` baseline extends the bulk-only model by adding a lightweight cell-type branch. For each patient $p$, we obtain a $K$-dimensional cell-type fraction vector $f_p \in \mathbb{R}^K$ from ReDeconv and encode it using a small MLP into a 256-dimensional embedding $h_{\mathrm{ct}}$. We then fuse $h_{\mathrm{ct}}$ with the bulk embedding $h_{\mathrm{b}}$ using one of the candidate fusion strategies (Concatenation, Sum, or Gated fusion) and feed the fused representation into the same survival head as in the bulk-only baseline. The fusion strategy and the cell-type branch hidden dimension were tuned per cohort within the grid search space defined in Table S3 while keeping the bulk encoder and survival head identical to those in the bulk-only baseline for fair comparison.

*scSurv baseline.*
We follow the official `scSurv` library for both training and inference. Specifically, bulk RNA-seq samples are first deconvoluted into patient-level cell-state mixture weights using the scSurv pipeline, and the inferred cell weights are then used as the input feature vector for downstream survival models. Following our unified evaluation protocol, the scSurv-derived cell weight vectors are respectively fed into Cox, DeepSurv, and DeepHit survival heads, while no bulk gene expression features are used.

All scSurv models are trained independently for each cancer cohort. Unless otherwise specified, scSurv-specific hyperparameters are set to the default values provided by the official implementation. For consistency with other baselines

**Received:** Date Month Year. **Revised received:** Date Month Year. **Accepted:** Date Month Year
© The Author(s) YEAR. Copyright and licence statements to be updated by the publisher during production.

| | Cohort statistics | | Reference scRNA-seq | | Gene statistics | | |
|---|---|---|---|---|---|---|---|
| **Cancer** | **Bulk** | **scRNA-seq** | **Cells** | **Cell types** | **Gene intersect** | **DEG** | **Folds** |
| COAD | 480 | 89 | 69,504 | 16 | 28,951 | 3,209–3,390 | 5 |
| BRCA | 1,095 | 25 | 99,290 | 9 | 17,929 | 2,612–2,831 | 5 |
| LUAD | 509 | 25 | 81,834 | 10 | 18,001 | 3,478–3,576 | 5 |
| LIHC | 372 | 6 | 21,504 | 6 | 17,157 | 2,732–2,875 | 5 |
| LGG | 516 | 14 | 20,995 | 6 | 17,882 | 3,451–3,533 | 3 |
| STAD | 408 | 26 | 112,718 | 11 | 18,169 | 2,763–3,082 | 5 |
| HNSC | 521 | 18 | 21,164 | 8 | 15,265 | 2,825–3,034 | 5 |
| KIRC | 535 | 7 | 17,188 | 7 | 15,421 | 3,648–3,897 | 5 |

**Table S1.** Comprehensive dataset statistics across cancer types. Cohort statistics summarize bulk RNA-seq survival samples and scRNA-seq donors. Reference scRNA-seq statistics report the total number of cells and annotated cell types. Gene statistics include the intersected gene sets between bulk RNA-seq and scRNA-seq and the min–max range of differentially expressed genes (DEGs) across cross-validation folds.

| Hyperparameter | Search Values |
|---|---|
| Hidden dim | $\{[512, 256], [1024, 512], [2048, 1024]\}$ |
| Batch normalization | {True, False} |
| Dropout | $\{0.4, 0.5\}$ |
| Activation | {ReLU, GELU, Tanh} |
| Batch size | $\{128, 256, 512\}$ |
| Learning rate | $\{10^{-2}, 10^{-3}, 10^{-4}\}$ |
| Weight decay | $\{10^{-2}, 10^{-3}\}$ |
| Alpha | $\{0.1, 0.2\}$ |
| Sigma | $\{0.1, 0.2\}$ |

**Table S2.** Grid search space for bulk-only survival models.

| Hyperparameter | Search Values |
|---|---|
| Batch size | $\{64, 128, 256, 512\}$ |
| Learning rate | $\{10^{-2}, 10^{-3}, 10^{-4}\}$ |
| Weight decay | $\{0, 10^{-2}, 10^{-3}, 10^{-4}\}$ |
| Dropout | $\{0.4, 0.5\}$ |
| Activation | {ReLU, GELU, Tanh} |
| Fusion strategy | {Concatenation, Sum, Gated} |
| Cell-type branch hidden dim | $\{64, 128, 256\}$ |

**Table S3.** Grid search space for the Bulk + Cell-type baseline.

| Hyperparameter | Search Values |
|---|---|
| VAE learning rate | $\{10^{-3}, 5 \times 10^{-3}, 10^{-2}\}$ |
| Deconvolution learning rate | $\{5 \times 10^{-3}, 10^{-2}, 5 \times 10^{-2}\}$ |
| Survival learning rate | $\{5 \times 10^{-5}, 10^{-4}, 5 \times 10^{-4}\}$ |
| Latent dimension | $\{32, 64\}$ |
| Hidden dimension | $\{128, 256\}$ |
| Hidden dims | $\{[64, 32], [128, 64], [256, 128]\}$ |
| Dropout | $\{0.4, 0.5\}$ |
| Activation | {ReLU, GELU, Tanh} |
| Alpha | $\{0.1, 0.2\}$ |
| Sigma | $\{0.1, 0.2\}$ |

**Table S4.** Grid search space for the scSurv baseline.

and DeSCENT, we use the same DEG-restricted gene set for deconvolution and survival modeling, instead of the highly variable genes (HVGs) adopted in the original scSurv paper. Hyperparameters associated with survival modeling are tuned per cohort via grid search, as summarized in Table S4.

| Hyperparameter | Search Values |
|---|---|
| Batch size | $\{16, 32, 64\}$ |
| Learning rate | $\{10^{-4}, 2 \times 10^{-4}, 5 \times 10^{-4}\}$ |
| Weight decay | $\{0, 10^{-3}, 10^{-2}\}$ |
| Dropout | $\{0.1, 0.2\}$ |
| Training epochs | $\{250, 300, 350\}$ |
| Embedding dimension | $\{256, 512\}$ |
| Number of attention heads | $\{4, 8\}$ |
| Number of layers | $\{1, 2\}$ |
| $\lambda_1$ (InfoNCE loss) | $\{0.5, 1.0, 2.0\}$ |
| $\lambda_2$ (Hard-negative loss) | $\{0.1, 0.5, 1.0\}$ |
| $\lambda_3$ (Mask reconstruction loss) | $\{0.1, 0.5, 1.0\}$ |
| Fusion strategy | {Cross-attention, Concat} |
| Pooling method | {Attention, Mean} |

**Table S5.** Grid search space for DeSCENT.

*DeSCENT.*

DeSCENT consists of a modality completion module and a multimodal representation module (main text, Section 3). The SCimilarity model was trained for 1000 epochs. The scDiffusion backbone was trained for 500 epochs with early stopping enabled and a patience of 100 epochs. For each patient, scDiffusion generated 2048 synthetic single cells. DeSCENT hyperparameters were tuned per cohort within the grid search space in Table S5.

| Cancer | ReDeconv | TAPE | Scaden | Ground Truth |
|---|---|---|---|---|
| COAD | **0.9902** | 0.9894 | 0.9882 | 0.9932 |
| BRCA | **0.9926** | 0.9683 | 0.9521 | 0.9976 |
| HNSC | **0.9968** | 0.9882 | 0.9860 | 0.9995 |
| KIRC | **0.9969** | 0.9964 | 0.9959 | 0.9980 |
| LGG | **0.9983** | 0.9839 | 0.9838 | 0.9992 |
| LIHC | 0.9888 | **0.9902** | 0.9890 | 0.9985 |
| LUAD | **0.9915** | 0.9911 | 0.9913 | 0.9990 |
| STAD | **0.9962** | 0.9853 | 0.9858 | 0.9981 |
| Average | **0.9939** | 0.9866 | 0.9840 | 0.9979 |

**Table S6.** Pearson correlation between generated and reference gene-expression profiles across cancers. **Bold** indicates the best deconvolution-based method, excluding Ground Truth.

| c.a. | m.r. | COAD | BRCA | LGG | LIHC | LUAD | STAD | HNSC | KIRC | Average |
|---|---|---|---|---|---|---|---|---|---|---|
| **DeSCENT - Cox** | | | | | | | | | | |
| ✗ | ✗ | $0.673_{0.030}$ | $0.706_{0.025}$ | $0.838_{0.019}$ | $0.699_{0.034}$ | $0.638_{0.019}$ | $0.590_{0.041}$ | $0.664_{0.024}$ | $0.728_{0.020}$ | 0.692 |
| ✗ | ✓ | $0.679_{0.052}$ | $\mathbf{0.729}_{0.040}$ | $0.843_{0.018}$ | $0.704_{0.025}$ | $0.645_{0.035}$ | $0.583_{0.043}$ | $0.654_{0.040}$ | $0.738_{0.019}$ | 0.697 |
| ✓ | ✗ | $0.681_{0.051}$ | $0.714_{0.020}$ | $\mathbf{0.847}_{0.025}$ | $0.705_{0.033}$ | $0.651_{0.070}$ | $0.604_{0.037}$ | $0.650_{0.039}$ | $0.734_{0.020}$ | 0.698 |
| ✓ | ✓ | $\mathbf{0.690}_{0.056}$ | $0.723_{0.012}$ | $0.845_{0.031}$ | $\mathbf{0.716}_{0.026}$ | $\mathbf{0.652}_{0.047}$ | $\mathbf{0.607}_{0.046}$ | $\mathbf{0.664}_{0.028}$ | $\mathbf{0.743}_{0.025}$ | **0.705** |
| **DeSCENT - DeepSurv** | | | | | | | | | | |
| ✗ | ✗ | $0.659_{0.033}$ | $0.688_{0.035}$ | $0.846_{0.034}$ | $0.700_{0.033}$ | $0.647_{0.037}$ | $0.602_{0.059}$ | $0.644_{0.060}$ | $0.738_{0.019}$ | 0.691 |
| ✗ | ✓ | $0.690_{0.049}$ | $0.700_{0.027}$ | $\mathbf{0.855}_{0.011}$ | $0.698_{0.050}$ | $0.677_{0.023}$ | $0.602_{0.044}$ | $\mathbf{0.659}_{0.041}$ | $0.734_{0.020}$ | 0.702 |
| ✓ | ✗ | $0.689_{0.042}$ | $0.711_{0.022}$ | $0.848_{0.026}$ | $0.709_{0.042}$ | $0.669_{0.028}$ | $0.610_{0.045}$ | $0.657_{0.040}$ | $\mathbf{0.743}_{0.025}$ | 0.704 |
| ✓ | ✓ | $\mathbf{0.708}_{0.038}$ | $\mathbf{0.723}_{0.019}$ | $0.853_{0.019}$ | $\mathbf{0.717}_{0.038}$ | $\mathbf{0.702}_{0.038}$ | $\mathbf{0.625}_{0.048}$ | $0.655_{0.030}$ | $0.732_{0.020}$ | **0.714** |
| **DeSCENT - DeepHit** | | | | | | | | | | |
| ✗ | ✗ | $0.651_{0.070}$ | $0.687_{0.042}$ | $0.818_{0.025}$ | $0.701_{0.037}$ | $0.645_{0.059}$ | $0.589_{0.042}$ | $0.650_{0.012}$ | $0.730_{0.019}$ | 0.684 |
| ✗ | ✓ | $0.673_{0.053}$ | $0.695_{0.025}$ | $0.825_{0.020}$ | $0.715_{0.029}$ | $0.653_{0.052}$ | $0.611_{0.045}$ | $0.655_{0.021}$ | $0.727_{0.018}$ | 0.694 |
| ✓ | ✗ | $0.687_{0.109}$ | $0.701_{0.015}$ | $0.820_{0.032}$ | $0.714_{0.030}$ | $0.651_{0.032}$ | $0.607_{0.047}$ | $0.659_{0.019}$ | $0.726_{0.022}$ | 0.696 |
| ✓ | ✓ | $\mathbf{0.696}_{0.076}$ | $\mathbf{0.703}_{0.038}$ | $\mathbf{0.826}_{0.035}$ | $\mathbf{0.731}_{0.020}$ | $\mathbf{0.670}_{0.018}$ | $\mathbf{0.627}_{0.038}$ | $\mathbf{0.660}_{0.007}$ | $\mathbf{0.736}_{0.024}$ | **0.706** |

**Table S9.** Ablation study on component effectiveness (C-index ± standard deviation). c.a.: contrastive alignment; m.r.: mask reconstruction.

| Cancer | ReDeconv | TAPE | Scaden | Ground Truth |
|---|---|---|---|---|
| COAD | **0.9775** | 0.9765 | 0.9724 | 0.9776 |
| BRCA | **0.9807** | 0.9587 | 0.9383 | 0.9895 |
| HNSC | **0.9921** | 0.9829 | 0.9806 | 0.9949 |
| KIRC | **0.9801** | 0.9783 | 0.9776 | 0.9818 |
| LGG | **0.9913** | 0.9773 | 0.9776 | 0.9929 |
| LIHC | 0.9863 | **0.9887** | 0.9880 | 0.9933 |
| LUAD | **0.9863** | 0.9790 | 0.9789 | 0.9903 |
| STAD | **0.9865** | 0.9805 | 0.9809 | 0.9901 |
| Average | **0.9851** | 0.9777 | 0.9743 | 0.9888 |

**Table S7.** Spearman rank correlation between generated and reference gene-expression profiles across cancers. **Bold** indicates the best deconvolution-based method, excluding Ground Truth.

| Cancer | ReDeconv | TAPE | Scaden | Ground Truth |
|---|---|---|---|---|
| COAD | **0.0330** | 0.0467 | 0.0463 | 0.0321 |
| BRCA | **0.0246** | 0.0493 | 0.0732 | 0.0151 |
| HNSC | **0.0122** | 0.0379 | 0.0436 | 0.0048 |
| KIRC | 0.0327 | 0.0278 | **0.0274** | 0.0260 |
| LGG | **0.0087** | 0.0569 | 0.0563 | 0.0086 |
| LIHC | 0.0356 | **0.0320** | 0.0350 | 0.0090 |
| LUAD | **0.0085** | 0.0342 | 0.0340 | 0.0036 |
| STAD | **0.0173** | 0.0762 | 0.0748 | 0.0148 |
| Average | **0.0216** | 0.0451 | 0.0488 | 0.0142 |

**Table S8.** Maximum Mean Discrepancy (MMD) between generated and reference gene-expression distributions across cancers. Lower is better. **Bold** indicates the best deconvolution-based method, excluding Ground Truth.

### S3. Deconvolution Results

In the tables above, ReDeconv/TAPE/Scaden/Ground Truth denote the source of conditioning fractions used for generation, while all reported Pearson/Spearman/MMD scores are computed between generated and reference gene-expression profiles.

To evaluate the accuracy of generated single-cell RNA-seq data, we compare gene-level expression distributions between real reference scRNA-seq $X_{\text{sc}}^{\text{ref}} \in \mathbb{R}^{n \times g}$ and generated scRNA-seq $X_{\text{sc}}^{\text{gen}} \in \mathbb{R}^{n \times g}$. Let $\mathbf{x} = \text{vec}(X_{\text{sc}}^{\text{ref}}) \in \mathbb{R}^N$ and $\mathbf{y} = \text{vec}(X_{\text{sc}}^{\text{gen}}) \in \mathbb{R}^N$ denote the flattened expression vectors, where $N = ng$.

*Pearson correlation.*

Pearson correlation measures the linear association between reference and generated expression values,

$$\text{Pearson} = \frac{\sum_{i=1}^{N}(x_i - \bar{x})(y_i - \bar{y})}{\sqrt{\sum_{i=1}^{N}(x_i - \bar{x})^2}\sqrt{\sum_{i=1}^{N}(y_i - \bar{y})^2}}, \tag{1}$$

where $x_i$ and $y_i$ denote the $i$-th elements of $\mathbf{x}$ and $\mathbf{y}$, respectively, and $\bar{x} = \frac{1}{N}\sum_{i=1}^{N} x_i$ (similarly for $\bar{y}$).

*Spearman correlation.*

Spearman correlation evaluates monotonic agreement and is computed as the Pearson correlation between ranked expression values,

$$\text{Spearman} = \text{Pearson}\big(\text{rank}(\mathbf{x}), \text{rank}(\mathbf{y})\big). \tag{2}$$

*Maximum Mean Discrepancy (MMD).*

To assess distributional similarity between reference and generated expression profiles, we compute the Maximum Mean Discrepancy (MMD),

$$\text{MMD}^2 = \left\| \mathbb{E}_{x \sim X_{\text{sc}}^{\text{ref}}}[\phi(x)] - \mathbb{E}_{y \sim X_{\text{sc}}^{\text{gen}}}[\phi(y)] \right\|_{\mathcal{H}}^2, \tag{3}$$

where $x$ and $y$ denote single-cell expression vectors sampled from the corresponding datasets, $\mathcal{H}$ is the reproducing kernel Hilbert space (RKHS) induced by a positive-definite kernel $k$, and $\phi(\cdot)$ is its associated feature map such that $k(a, b) = \langle \phi(a), \phi(b) \rangle_{\mathcal{H}}$. We use a Gaussian kernel $k(a, b) = \exp(-\|a - b\|^2/2\sigma^2)$, with bandwidth $\sigma$ selected via the median heuristic. Lower MMD values indicate closer agreement between reference and generated expression distributions.

### S4. Ablation Study

We investigated the effects of contrastive alignment and mask reconstruction by toggling these losses in our training objective. The two contrastive terms, InfoNCE and hard negative, are enabled or disabled together. Results across eight cancer types and three survival heads are reported in the supplementary material. Enabling both contrastive alignment and mask reconstruction achieves the best average performance, indicating that patient-level cross-modal semantic alignment and gene-level self-supervised regularization provide complementary benefits for learning robust fused representations. Even when both objectives are disabled, DeSCENT maintains a non-trivial improvement over bulk-only models, suggesting that multimodal fusion with patient-specific synthetic single-cell inputs already contributes substantially to performance gains, while the auxiliary objectives further amplify these benefits.
